# Rapid estimation of insect biodiversity using AI-assisted image analysis: a cost-effective tool for monitoring and ecological assessment

**DOI:** 10.64898/2026.11.11.765925

**Authors:** Liraz Cabra-Leykin, Tamar Keasar, Chen Keasar

**Affiliations:** Department of Evolutionary and Environmental Biology, University of Haifa, Israel; Department of Biology, University of Haifa – Oranim, Israel; Faculty of Computer and Information Science, Ben-Gurion University of the Negev, Israel

**Keywords:** bioindicators, conservation, ecological assessment, ecosystem services, insect biodiversity, image processing, machine learning

## Abstract

Protection of natural areas and environmentally friendly management of anthropogenic landscapes require reliable tools for monitoring and comparing biodiversity. Bioindicator-based approaches can support large-scale ecological assessment, but insect monitoring remains constrained by the taxonomic expertise required for species identification. Here, we propose an AI-supported index for estimating the biodiversity of flying insects, based on the diversity of visual features of insect bodies. Flying insects were captured using yellow sticky traps across agricultural and natural habitats, seasons, crops, and sites in Israel’s Mediterranean region. Traps were scanned at high resolution and uploaded to a dedicated image database. A machine learning model was used to detect and delineate insects, after which engineered visual features representing body size, color, and shape were extracted. Biodiversity was estimated from distances among feature vectors and compared with taxonomic diversity derived from expert identification of a representative validation set. The AI-derived diversity index correlated strongly with expert-based taxonomic diversity, supporting the hypothesis that morphological variation in insect images can provide a proxy for taxonomic diversity. Applying the index to the full dataset revealed a negative relationship between insect abundance and diversity, while diversity also varied across habitats and sampling seasons. Our results demonstrate the potential of AI-based image analysis to provide rapid and cost-effective assessments of insect biodiversity without requiring complete taxonomic identification. This approach may facilitate biodiversity monitoring and support timely ecological assessment and decision-making in sustainable agricultural and landscape management.

## 1. Introduction

### The challenge

Biodiversity is declining at an alarming rate due to human activities. This poses a significant threat to the environment, human health, and the stability of ecosystems. Habitat degradation and loss are the main drivers of the biodiversity crisis, mainly due to land conversion to agriculture, mines, energy production facilities, cities, and roads (IPBES, 2019). Protection of natural areas and environmentally friendly management of anthropogenic landscapes are key strategies for biodiversity conservation. Prioritizing conservation efforts and assessing the effectiveness of such interventions is often challenging, especially in light of accelerating development processes. This generates an increasing need for comparative tools to monitor and assess biodiversity in a reliable, rapid, and cost-effective manner (Galler et al., 2015; Long et al., 2018; Mörtberg et al., 2007).

### Insects as indicators

Monitoring a suite of bioindicators is a popular approach to large-scale assessment of the quality and health of ecosystems (Burger, 2006; Stephenson, 2020). Arthropod higher taxa were found to be good biotic indicators of ecosystem services and dis-services because of their ease of sampling and diverse ecological roles. Insects and other arthropods play crucial beneficial roles in crop pollination and biological control, but also act as crop pests and disease vectors. Drastic changes in arthropod population abundances and diversity have negative cascading effects on ecological stability and ecosystem resilience (Billeter et al., 2007; Duelli and Obrist, 2003; Kent and Smart, 1981; Wathern et al., 1986).

### AI for insect monitoring

Insect assemblages typically include numerous, small-bodied, individuals of diverse taxa. Thus, their composition is routinely inferred from trapping-based samples. Traditionally, the caught specimens are classified by experts, based on morphological characteristics, and the taxonomic counts are statistically analysed. This workflow is constrained by extreme labor intensity, high logistical costs, and a global shortage of taxonomists capable of identifying the world’s vast insect diversity (Gao et al., 2024; Kalfas et al., 2023; Preti et al., 2021; Schneider et al., 2022). This systematic disparity between urgent needs and low availability calls for technological solutions that can ease the need for taxonomic expertise. Current approaches include: genetic procedures, such as DNA barcoding (J. S. et al., 2024) and metabarcoding (Buchner et al., 2025); sensors such as radar (Tielens et al., 2025), on-trap microphones (Seyidbayli et al., 2025), and cameras combined with image processing (Biswas and Tiwari, 2024; Geissmann et al., 2022; Szczygieł et al., 2026a). A comprehensive review of these solutions is beyond the scope of this manuscript and the reader is referred to a recent one (Van Klink et al., 2024).

### The knowledge gap in existing AI-supported methods

Most of the current deep learning models in entomology were developed for Integrated Pest Management (IPM). The economic necessity of protecting global food supplies has made pest control the dominant driver for the development and deployment of these technologies (Chiavassa et al., 2024; Chiranjeevi et al., 2024; Gao et al., 2024; Kittichai et al., 2021). Thus, many deep learning models focus on identifying and monitoring insects that harm agricultural crops (Chiranjeevi et al., 2024; Ciampi et al., 2023; Teixeira et al., 2023). Related models aim to monitor public health threats, such as mosquito vectors of disease (Kittichai et al., 2021). Nevertheless, biodiversity assessment is a significant and growing field in AI-assisted entomology.

Biodiversity assessment campaigns differ from pest monitoring projects in two important aspects. First, they should ideally use low-cost trapping methods to allow broad sampling of insect communities over space and time. In this respect, low-tech methods such as pitfall traps and sticky traps are more suitable than camera traps, which are often used to monitor crop pests and disease vectors (Bjerge et al., 2023; Colares et al., 2025; Fujisawa et al., 2023; Schneider et al., 2022). In addition, AI-assisted assessment of biodiversity must cope with the vast diversity of insect species, whereas pest monitoring efforts focus on a few well-known target species. Many of the world’s insect species, especially in the global south, are undescribed or cryptic (i.e., can be identified genetically but not morphologically). Even when morphological identification is possible, large-scale standardized data for training machine learning models is often lacking for rare species. This challenge has so far been addressed by estimating insect biodiversity at broader taxonomic levels, such as families (Bjerge et al., 2023) or orders (Colares et al., 2025). Yet even coarse classification tasks (e.g., assigning insects to one of ∼20 orders) require extensive data collection efforts and development of public, curated reference databases for model training. These challenges motivate the search for biodiversity proxies that can be rapidly estimated from large-scale datasets of trapped insects even when training data are scarce.

### Our approach

We focus on combining a low-tech field method - capturing insects using sticky traps - with automated analysis via image processing of the scanned traps. (Fig. 1). This approach is arguably highly cost effective, and thus scales well and allows comprehensive insect surveys over wide spatial and temporal ranges. Sticky traps passively catch airborne arthropods, mainly insects but also ballooning spiders. They have been used by entomologists for decades as a simple and cheap means of monitoring flying insects. The traps are placed in the field for a few hours or days, and are then inspected by experts. While manual inspection of insect-loaded sticky traps is still the mainstream approach, recent years have also witnessed the emergence of computerised, on-trap camera systems. Notwithstanding their important contributions, trap-camera systems weaken two of the main advantages of sticky traps: low price and resilience to harsh environmental conditions. We have recently described an alternative approach for sticky trap digitization, scanning them using standard office scanners (Keasar et al., 2024). We have also created a database of scanned sticky trap images (STARdbi). Here we propose an approach towards automatic biodiversity estimation of the scanned trap images. Under the working hypothesis that taxonomic diversity is manifested by visual diversity, we extract visual features from images of trapped insects using image processing tools (section 2.3) and represent them in multidimensional feature spaces. To this end, we use biological intuition to develop engineered features, related to the insects’ size, shape, and color distributions (Methods, section 2.3.2). A visual diversity index (VDI) is calculated from the shape of the resulting point clouds.

**Figure 1:**
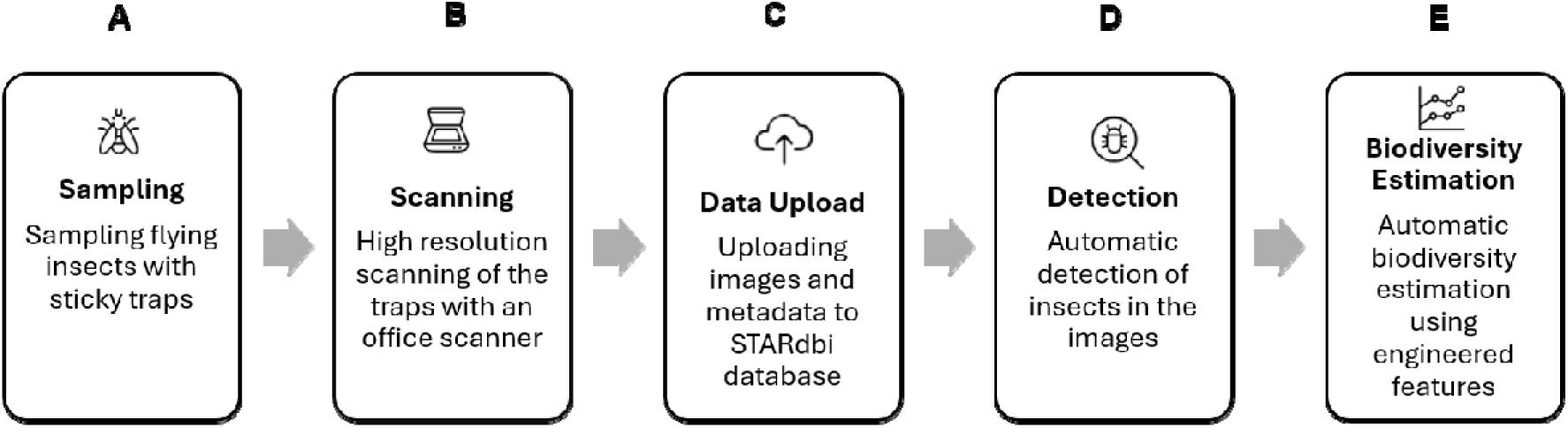
Automatic insect detection and biodiversity estimation workflow.

Making sense of a set of observations, represented as a point cloud in a (typically) multidimensional space, is a common task in data science. It often relies on estimating an underlying probability density function from which the observations were sampled, or at least some of its properties. Often, the points cannot be adequately described by standard parametric distributions (e.g., Gaussian), calling for nonparametric methods. Some methods consider the discreteness of the sample by replacing each point by a smoothly decaying function, known as “kernel” (Epanechnikov, 1969; Parzen, 1962; Rosenblatt, 1956). In this study, we represent individual arthropods as points in a multidimensional space of visual features. An assemblage of individuals (e.g., arthropods caught on a single trap) is a sample of the environment’s community, and its visual diversity may serve as a proxy for the taxonomic diversity of the community (here estimated by the Shannon-Weaver index). We have experimented with several methods to estimate this diversity, and converged to a grid-based set of pseudo-species, discussed below. We dub this method Grid Pseudo-Species Visual Diversity Index (GPS-VDI). We use a Gaussian kernel to assign individuals to these pseudo-species. This is similar in spirit to soft (or fuzzy) histograms used in image processing (Plant et al., 2012; Reznik et al., 2010). In ecological context, a similar remote-sensing-based approach has recently been proposed for plant diversity (Pinto-Ledezma et al., 2025).

### Aims

Our research aims to develop an AI-supported index estimating the biodiversity of flying insects as bioindicators combined with a low-cost field insect monitoring method using sticky traps. We compare our computational method for classifying insects to the traditional method by experts, with the hope that this field will develop in the coming years and lead to a breakthrough in entomological research. An AI-supported index allows rapid estimation of arthropod biodiversity and provides a tool for ecological assessment using insects as bioindicators, which can address the need for timely decision-making in sustainable landscape planning and management.

## 2. Materials and methods

### 2.1 Study area

We sampled 30 sites along a ∼200 km north-south topographic and climatic gradient that spanned a-200 - 1050 elevation range and a 250-900 mm annual precipitation range. The sites represent ecological units and land-use categories in the open landscape of Israel. In 11 sites, we sampled two agricultural plots of the same crop type and one nearby natural plot; these plots were sampled repeatedly across different seasons and years, in 3-4 sampling rounds. In 19 sites, only two agricultural plots were sampled during a single round in the spring. In total, there were six sampling rounds: in August 2022, May 2023, May 2024, August 2024, May 2025 and August 2025. This intensive sampling allowed us to compare the diversity of the trapped insects between habitats and seasons. This design resulted in a total of 69 sampling plots (Table 1). The agricultural plots are characterized by intensive cultivation and represent major crops: Rosaceae (apple, pear, nectarine, plum, cherry, and almond), citrus (grapefruit, pomelo, orange, and lemon), Evergreen (avocado, lychee, mango, olive), grape vineyards and annual crops. The natural habitat is characterized by Mediterranean herbs and shrubs with minimal disturbance, typical of northern Israel.

**Table 1.** Study sites and sampling plots list. The plots highlighted in grey were selected for the paired-plot test in Section 2.6 of the Methods.

| Plot id | Site | Habitat | Crop | Major crops | Altitude (m) | Rainfall - annual mean (mm) | X | Y |
|---|---|---|---|---|---|---|---|---|
| 1 | Afik | Agriculture | Pear | Deciduous | 346 | 487 | 266575 | 742184 |
| 2 | Afik | Agriculture | Pear | Deciduous | 344 | 488 | 266743 | 742032 |
| 3 | Afik | Natural |  |  | 343 | 485 | 266299 | 742049 |
| 4 | Naot Golan | Agriculture | Almond | Deciduous | 379 | 506 | 268416 | 744249 |
| 5 | Naot Golan | Agriculture | Almond | Deciduous | 375 | 506 | 268519 | 744256 |
| 6 | Naot Golan | Natural |  |  | 370 | 507 | 268753 | 744210 |
| 7 | Nov South | Agriculture | nectarine | Deciduous | 385 | 524 | 272550 | 745651 |
| 8 | Nov South | Agriculture | nectarine | Deciduous | 384 | 524 | 272685 | 745621 |
| 9 | Nov South | Natural |  |  | 379 | 522 | 272540 | 744991 |
| 10 | Nov North | Agriculture | Plum | Deciduous | 387 | 526 | 272388 | 746232 |
| 11 | Nov North | Agriculture | Plum | Deciduous | 388 | 525 | 272394 | 745988 |
| 12 | Nov North | Natural |  |  | 380 | 526 | 272282 | 746531 |
| 13 | Yonatan | Agriculture | Cherry | Deciduous | 651 | 657 | 277536 | 761236 |
| 14 | Yonatan | Agriculture | Cherry | Deciduous | 646 | 657 | 277467 | 761216 |
| 15 | Yonatan | Natural |  |  | 664 | 660 | 277749 | 761451 |
| 16 | Merom Golan | Agriculture | Apple | Deciduous | 942 | 831 | 274834 | 781946 |
| 17 | Merom Golan | Agriculture | Apple | Deciduous | 943 | 829 | 274465 | 782018 |
| 18 | Merom Golan | Natural |  |  | 987 | 813 | 274446 | 781129 |
| 19 | Elrom West | Agriculture | Apple | Deciduous | 1031 | 919 | 274967 | 786231 |
| 20 | Elrom West | Agriculture | Apple | Deciduous | 1043 | 924 | 274952 | 786512 |
| 21 | Elrom West | Natural |  |  | 1021 | 912 | 274700 | 785949 |
| 22 | Elrom East | Agriculture | Apple | Deciduous | 1032 | 920 | 275273 | 786314 |
| 23 | Elrom East | Agriculture | Apple | Deciduous | 1021 | 921 | 275496 | 786221 |
| 24 | Elrom East | Natural |  |  | 1019 | 923 | 275630 | 786122 |
| 25 | Mevo Hama | Agriculture | Almond | Deciduous | 344 | 426 | 262202 | 737771 |
| 26 | Mevo Hama | Agriculture | Almond | Deciduous | 341 | 427 | 262359 | 738022 |
| 27 | Mevo Hama | Natural |  |  | 358 | 406 | 261345 | 736721 |
| 28 | Kidmat Tzvi | Agriculture | Pear | Deciduous | 503 | 594 | 265716 | 772142 |
| 29 | Kidmat Tzvi | Agriculture | Pear | Deciduous | 509 | 594 | 265716 | 772142 |
| 30 | Kidmat Tzvi | Natural |  |  | 518 | 605 | 265847 | 772177 |
| 31 | Maale Gilboa | Agriculture | Grape | Grape | -177 | 283 | 247651 | 701634 |
| 32 | Maale Gilboa | Agriculture | Grape | Grape | -177 | 283 | 247888 | 700893 |
| 33 | Maale Gilboa | Natural |  |  | -166 | 287 | 247185 | 702104 |
| 34 | Moledet | Agriculture | Oroblanco | Citrus | 45 | 405 | 242183 | 720677 |
| 35 | Moledet | Agriculture | Oroblanco | Citrus | 41 | 404 | 242060 | 720630 |
| 36 | Hukok | Agriculture | Avocado | Evergreen | -71 | 494 | 246811 | 753226 |
| 37 | Hukok | Agriculture | Avocado | Evergreen | -54 | 491 | 246903 | 753366 |
| 38 | Ravid | Agriculture | Avocado | Evergreen | 104 | 509 | 244104 | 750388 |
| 39 | Ravid | Agriculture | Avocado | Evergreen | 104 | 504 | 244196 | 750376 |
| 40 | Beit Zera | Agriculture | Avocado | Evergreen | -204 | 391 | 253849 | 731954 |
| 41 | Beit Zera | Agriculture | Avocado | Evergreen | -204 | 391 | 253705 | 731977 |
| 42 | Kineret East | Agriculture | Avocado | Evergreen | 160 | 426 | 249203 | 741105 |
| 43 | Kineret East | Agriculture | Avocado | Evergreen | 165 | 427 | 249317 | 741214 |
| 44 | Kineret West | Agriculture | Almond | Deciduous | 149 | 428 | 249035 | 741049 |
| 45 | Kineret West | Agriculture | Almond | Deciduous | 138 | 428 | 248885 | 740989 |
| 46 | Aniam | Agriculture | Grapefruit | Citrus | 55 | 510 | 259245 | 762032 |
| 47 | Aniam | Agriculture | Grapefruit | Citrus | 52 | 510 | 259235 | 762196 |
| 48 | Mop Hula Valley | Agriculture | Avocado | Evergreen | 74 | 467 | 258660 | 784152 |
| 49 | Mop Hula Valley | Agriculture | Avocado | Evergreen | 71 | 464 | 258672 | 784043 |
| 50 | Gaaton | Agriculture | Avocado | Evergreen | 35 | 624 | 212926 | 765458 |
| 51 | Gaaton | Agriculture | Avocado | Evergreen | 37 | 624 | 213029 | 765402 |
| 52 | Gimzo | Agriculture | Clementine | Citrus | 95 | 550 | 193843 | 648297 |
| 53 | Gimzo | Agriculture | Clementine | Citrus | 90 | 550 | 193775 | 648212 |
| 54 | Naan | Agriculture | Clementine | Citrus | 82 | 550 | 187130 | 643462 |
| 55 | Naan | Agriculture | Clementine | Citrus | 88 | 549 | 187197 | 643560 |
| 56 | Hulda | Agriculture | Olive | Evergreen | 103 | 537 | 188369 | 638547 |
| 57 | Hulda | Agriculture | Olive | Evergreen | 97 | 537 | 188466 | 638717 |
| 58 | Bekoa | Agriculture | Orange | Citrus | 138 | 529 | 194258 | 636868 |
| 59 | Bekoa | Agriculture | Orange | Citrus | 141 | 529 | 194182 | 636782 |
| 60 | Revadim | Agriculture | Avocado | Evergreen | 81 | 522 | 182436 | 630674 |
| 61 | Revadim | Agriculture | Avocado | Evergreen | 80 | 522 | 182382 | 630644 |
| 62 | Tlamim West | Agriculture | Clementine | Citrus | 74 | 432 | 167602 | 607138 |
| 63 | Tlamim West | Agriculture | Clementine | Citrus | 74 | 434 | 167544 | 607187 |
| 64 | Tlamim East | Agriculture | Lemon | Citrus | 76 | 433 | 167855 | 607235 |
| 65 | Tlamim East | Agriculture | Lemon | Citrus | 76 | 433 | 167860 | 607144 |
| 66 | Klahim | Agriculture | Cabbage | Annual | 179 | 330 | 170625 | 595881 |
| 67 | Klahim | Agriculture | Cabbage | Annual | 174 | 329 | 170711 | 595724 |
| 68 | Maale Gilboa - APV1 | Agriculture | Lychee | Evergreen | -173 | 285 | 247412 | 701938 |
| 69 | Maale Gilboa - APV2 | Agriculture | Mango | Evergreen | -172 | 285 | 247499 | 701868 |

### 2.2 Insect sampling

We used yellow sticky traps to monitor flying insects (Fig. 1A). We set up three 10×25 cm traps in each plot and sampling round, totaling 327 traps. Five of these traps disappeared during the deployment in the field and could not be analyzed. The traps were mounted on a tree or a pole at a height of approximately 1.5 m, and were positioned ∼10 m inside the plot boundary to minimize edge effects. Each trap was exposed with a single adhesive surface facing outward for 10 days, after which it was covered with a transparent acetate sheet to preserve specimen integrity during transport. Traps were subsequently digitized (Fig. 1B) for the subsequent image analyses (Fig. 1C-E), as detailed in section 2.3 below. The scanned traps are deposited in the Margolin House Natural History collections at Oranim College.

### 2.3 Image processing

The raw data for this study consist of sticky trap RGB images. The acetate-coated traps were scanned in high resolution (1,200 dpi), using standard office scanners (Fig.1B). The images were uploaded to the STARdbi database (Keasar et al., 2024) along with their metadata (Fig. 1C). The insects on the traps were detected (section 2.3.1, Fig. 1D). The biodiversity of each trap was then estimated from its set of caught insects based on five visual features (section 2.3.2) (Fig. 1E).

#### 2.3.1 Automatic insect detection (Fig. 1D)

The sticky traps are conceptually considered as samples of the flying insect community in a given place and time. In practice, however, the individual insects occupy a small fraction of the trap surface and are accompanied by irrelevant wind-carried debris. Thus, any visual inference from the trap scans requires the extraction of insect crops, while keeping their association with the trap metadata: location, times of placement and removal, etc. From a computational perspective, this is a special case of object detection, a standard computer vision task (Zou et al., 2023).

To address this task, we have assembled a dataset of 416 traps from numerous ongoing research projects, for which areas-of-interest (AoIs - boxes that enclose insects) were manually assigned, using the user interface provided by STARdbi. Eighty-four traps (18,893 individual arthropods) were set aside as a test set and the remaining 332 traps (51,458 arthropods) were used as a training set.

The traps are deliberately much larger than individual insects, to allow the capture of many of them. The resulting orders of magnitude size difference between the insects and the traps complicates the naive use of off-the-shelf methods. Specifically, the trap scans are at least 10M pixels in size. Standard packages for object detection downscale such large images by at least an order of magnitude, to accommodate memory constraints. In order to avoid the loss of minute insects (often less than 200 pixels) we split the images to overlapping 2000×2000 pixel tiles. These tiles were augmented (three 90-degree rotations) and used to train a Faster-RCNN (Ren et al., 2015) object detection model (detectron2 implementation (Wu et al., 2019). In the inference stage, tile annotations are merged to provide trap-based annotation, taking care of duplicated annotations.

#### 2.3.2 Engineered features (Fig. 1E)

Our basic unit of analysis is an image containing a single arthropod, extracted from the trap image. Aside from the arthropod itself, these images contain only the originally uniform trap background, to which field exposure introduces some variation due to dust and debris. Thus, variation among images may be attributed primarily to variation in the visual characteristics of the arthropods. Specifically, we use five visual features of three different types, to represent each arthropod image:

1. Arthropod size (log-transformed) - The projected size of each arthropod was estimated by separating it from the trap background. Briefly, each detected arthropod was cropped from the image. The typical background color and its variation were estimated from the parent trap image, and pixels sufficiently different from this background were provisionally classified as belonging to the arthropod. Smoothing and morphological operations were then used to remove small artifacts and consolidate the resulting mask. The number of pixels in the final mask, normalized to a scanning resolution of 1200 dpi, was taken as the arthropod’s projected size. Because arthropod sizes span a wide range, their logarithms were used as the visual feature.
2. Insect shape (e.g., rounded vs. elongated) - the arthropod’s color image is first converted to grayscale and decomposed using singular value decomposition (SVD), to generate a vector (⊠_0_,⊠_1_,⊠_2_,…,⊠_⊠1_)of singular values ordered from larger to smaller, which capture progressively weaker components of the image spatial intensity pattern. Simple, regular images tend to be dominated by a few components, whereas more complex intensity patterns are distributed among more components. We use 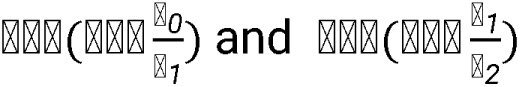 as two features that describe coarse aspects of arthropod shapes. While the ratios themselves are also strong features, they may occasionally reach very high values, compromising the method’s numerical stability.
3. Insect color - In an RGB image, each pixel is represented by a triplet of values (r, g, b), corresponding to its red, green, and blue intensities. We use the ratio of the total red intensity to the total green intensity, 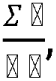 and the corresponding red-to-blue ratio, 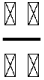 as two features that capture aspects of arthropod color. Notably, these descriptors consider all the image pixels including the background. Since the background is approximately identical in all the images, its contribution is expected to primarily reduce rather than generate variation among the color descriptors.

#### 2.3.3 Visual diversity estimation by GPS-VDI

Given ⊠ features, an arthropod assemblage, say those caught on a single trap, is represented by an ⊠ feature matrix ⊠, where each of the rows represent an individual, and each column represents one of the features. A preliminary analysis showed that the estimated diversity is sensitive to sample size. To cope with this bias, we calculate the diversity of a small sample of ⊠ rows (sampled without repetition), and report the average of a large number, ⊠, of such samples. Specifically, to calculate the diversity of the insect assemblage we:

1. Linearly scale the values into the [0, 1] range. The scaling is done per-feature using the minimum and maximum values in the union of all the studied assemblages, say all the traps in a given survey.
2. The-dimensional unit hypercube, which contains all the observations, is subdivided by evenly distributed positions along each dimension. This generates a grid, which consists of vertices. Each actual observed individual is mapped to vertices in its neighborhood. Those vertices that are associated with individuals may be loosely interpreted as pseudo-species, allowing the application of standard ecological indices to estimate their diversity. The user defined parameter controls the resolution of the feature space sampling. The larger, the finer is the sampling, but at the same time the computational price tag increases.
3. Formally, given⊠ ⊂ ⊠, a random sample of ⊠ rows (individuals) from matrix F, each pseudo-species (grid vertex), ⊠, is associated with an abundance score. This score is the distance-weighted contribution of all the sample’s arthropods to the vertex av = ∑sɛs K(v,s) where ⊠(⊠,⊠) = ⊠-^⊠⊠(⊠⊠)2^ where is a Gaussian kernel function, and ⊠(⊠,⊠) is the Euclidean distance between ⊠ and ⊠. The user defined parameter λ controls the effective neighborhood of individual arthropods. The smaller it is, the wider is this neighborhood, and thus the number of contributed pseud-species is larger. Notably, extreme values of λ (relative to the vertices spacing) render the⊠⊠values meaningless. A very small value will associate all the individuals with all the vertices, resulting in high and even abundance. Very high values will leave almost all the vertices empty, with a few, arbitrary, occupied ones.
4. The grid vertex ⊠ is then associated with normalised abundance 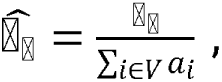 which is analogous to the relative species abundance used in ecology.
5. The visual diversity of the sample is then calculated as 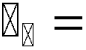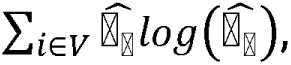 similarly to the calculation of the Shannon-Weaver index of taxonomic diversity.
6. Steps 1-4 are repeated L times, and the overall diversity is estimated as 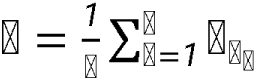
7. In the current study⊠ = 5, ⊠ = 5, λ = 40, ⊠ = 50, and ⊠ = 1000.

Data were analyzed using Python, with NumPy (Harris et al., 2020) and pandas (McKinney, 2010) used for data processing and analysis. Data were visualized using Matplotlib (Hunter, 2007), and kernel density estimates were generated using the kdeplot function in seaborn (Waskom, 2021).

#### 2.3.4 Sensitivity tests

As discussed below, the correlation between taxonomic and visual diversities of arthropod samples is the major observation of this study. While the actual numbers reported depend on the exact settings, the correlation itself needs to be qualitatively stable to minor perturbations. To validate this, we performed several tests:

1. Testing a range of grid resolution (*K*), and effective neighborhood size (λ)combinations.
2. Replacing the grid-based estimation of visual diversity with the more conventional KDE and KDE_loo methods (Epanechnikov, 1969; Parzen, 1962; Rosenblatt, 1956), using the KDE implementation of SkLearn (Gramfort et al., 2011).
3. Applying subsets of the visual features.
4. Generating synthetic assemblages where taxonomic groups come from different original traps. The Shannon-Weaver indices of these assemblages differ from those of the original traps, but they keep the visual-cues/taxon association.

### 2.4 Validation

To validate the automated insect diversity estimates, a manual taxonomic reference dataset was constructed by expert entomologists. A subset of 40 sticky traps representative of different land uses, crop types, and geographic regions was selected for expert annotation. All delineated insect detections from these traps were exported as individual image files for taxonomic identification, a total of 10,303 insect images. Insect images were classified across five hierarchical taxonomic levels: order, suborder, infraorder, superfamily, and family. The achievable level of classification varied and depended on the taxonomists’ expertise, insect preservation state, and image quality. In the first stage, images were sorted into major observed orders, including Diptera (flies), Coleoptera (beetles), Hemiptera (true bugs), Hymenoptera (bees, wasps and ants), Lepidoptera (butterflies and moths), Thysanoptera (thrips), Neuroptera (net-winged insects), and an “unidentified insect” category. High-quality insect images were then sorted to the family level, with intermediate sorting to the suborder, Infraorder and superfamily levels (Fig. 2). All taxa were resolved to identifiers using the Integrated Taxonomic Information System (ITIS TSN) (Supplementary Table S1). A total of 8,200 insect images were classified to the order level and a subset of 3,704 images were further classified to the family level. Two professional entomologists divided the dataset equally for initial sorting and then independently cross-checked each other’s classifications to ensure consistency and minimize potential observer bias. Each expert worked 2-4 hours on each trap, depending on the number of specimens caught.

**Figure 2:**
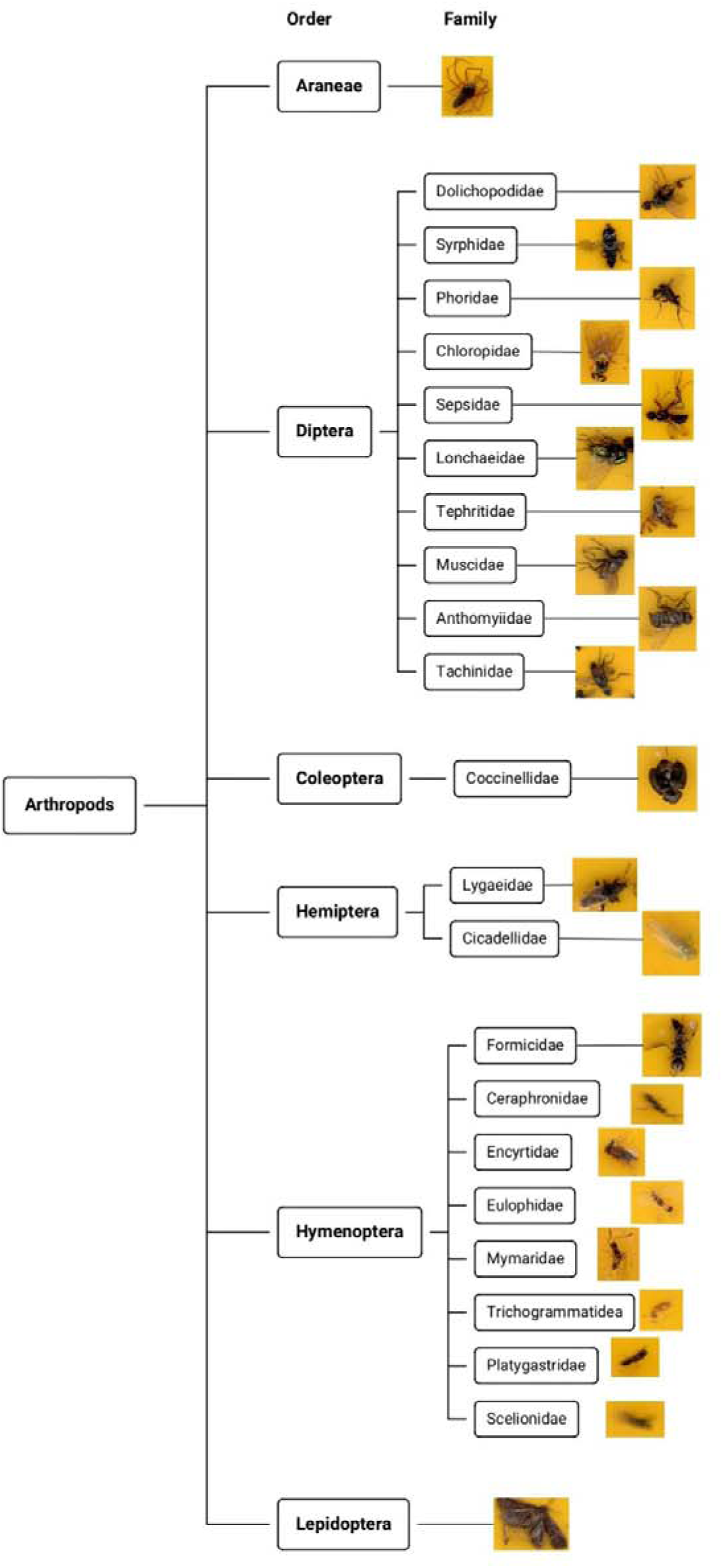
Classification of insects orders and families.

### 2.5 Data Analysis – Biodiversity: Computational Method vs. Taxonomic Experts

Order-level and family-level Shannon-Weaver diversity indices were calculated for each trap based on the expert-derived taxonomic classifications, representing the benchmark for biological diversity. The entropy-based diversity index was computed from the visual feature space of the insect images (section 2.3.2), reflecting phenotypic heterogeneity rather than taxonomic composition. To evaluate concordance between the two methods, we calculated Pearson’s correlation coefficients between the expert-derived taxonomic diversity values and those obtained from the visual-entropy approach, using both unweighted and weighted correlations. The unweighted (standard) correlation assigns equal weight to all traps, regardless of their occupancy. The weighted correlation assigns higher weight to dense traps than to sparse ones to further reduce sample size bias.

### 2.6 An illustrative field application: the effects of season and land use on the visual diversity of flying insects

We applied the ‘grid’ visual diversity estimator to the full dataset of 322 sticky traps. The resulting per-trap diversity estimates were then used to evaluate three ecological questions: (1) Is insect diversity related to insect abundance per trap? (2) Do insect diversity and abundance differ between natural and agricultural habitats? and (3) Do insect diversity and abundance differ between the spring and summer sampling seasons?

To facilitate comparisons among traps, insect diversity values were transformed to Z-scores prior to statistical analyses. The mean of the z-transformed distribution is zero. Positive and negative z-scores correspond to above-average and below-average diversity, respectively. The relationship between insect abundance (number of insects per sticky trap) and standardized insect diversity was assessed using Pearson’s correlation. To evaluate the effects of habitat and season on insect abundance and diversity, we used only 137 traps from the complete dataset, which comprised paired observations. These traps represent 11 sites, where traps were set up in triplicates in two habitats (agricultural and natural) and in two seasons (spring and summer). Nine sites were sampled over one year, and two were sampled over two years. Restricting the analysis to paired traps enabled direct comparisons between natural and agricultural habitats while minimizing spatial variation among sampling sites. Because of the repeated sampling design, and the non-independence of traps from each of the sampling sites, we fitted generalized linear mixed models (GLMMs) to the data. Habitat (natural vs. agricultural), season (spring vs. summer), and their interaction were included as fixed effects and site identity was treated as a random effect. Including year as a fixed effect did not improve model fit, and year was therefore omitted from the models. Separate models were fitted for insect abundance (number of insects per trap) and insect diversity. Diversity was analyzed using a gaussian distribution with an identity link function and abundance was analyzed using a poisson error distribution with a log link function. The models’ intercept represents a theoretical ‘baseline’ abundance and diversity in the study sites, without the contribution of habitat and season. Model significance was assessed using likelihood ratio tests, and pairwise post-hoc comparisons were conducted when appropriate. These analyses were also performed on triplicates (for the total insects in the three traps/repeats combined), allowing the three replicates to be treated as one larger trap. These results were also reported. Statistical analyses were conducted in R version 4.2.2 (R Core Team, 2022). Linear mixed-effects models were fitted using the lme4 package (Bates et al., 2015), with significance tests and p-values obtained using lmerTest (Kuznetsova et al., 2017).

## 3. Results

### 3.1 The hyperspace of visual features

Each of the captured insects is characterized by a vector of values that represent the five engineered visual features. This vector defines each individual’s coordinates in a five-dimensional hyperspace of features. A two-dimensional projection of the hyperspace demonstrates that taxonomic families cluster in distinct parts of its volume (Fig. 3). This is illustrated for four well-represented families within the expert-identified subset of images: leafhoppers (Cicadellidae), phorid flies (Phoridae), moth flies (Psychodidae), and fairy wasps (Mymaridae). We also note that some taxa (e.g., the fairy wasps) are more visually diverse than others (e.g., the moth flies), hence occupy a larger fraction of the hyperspace.

**Figure 3:**
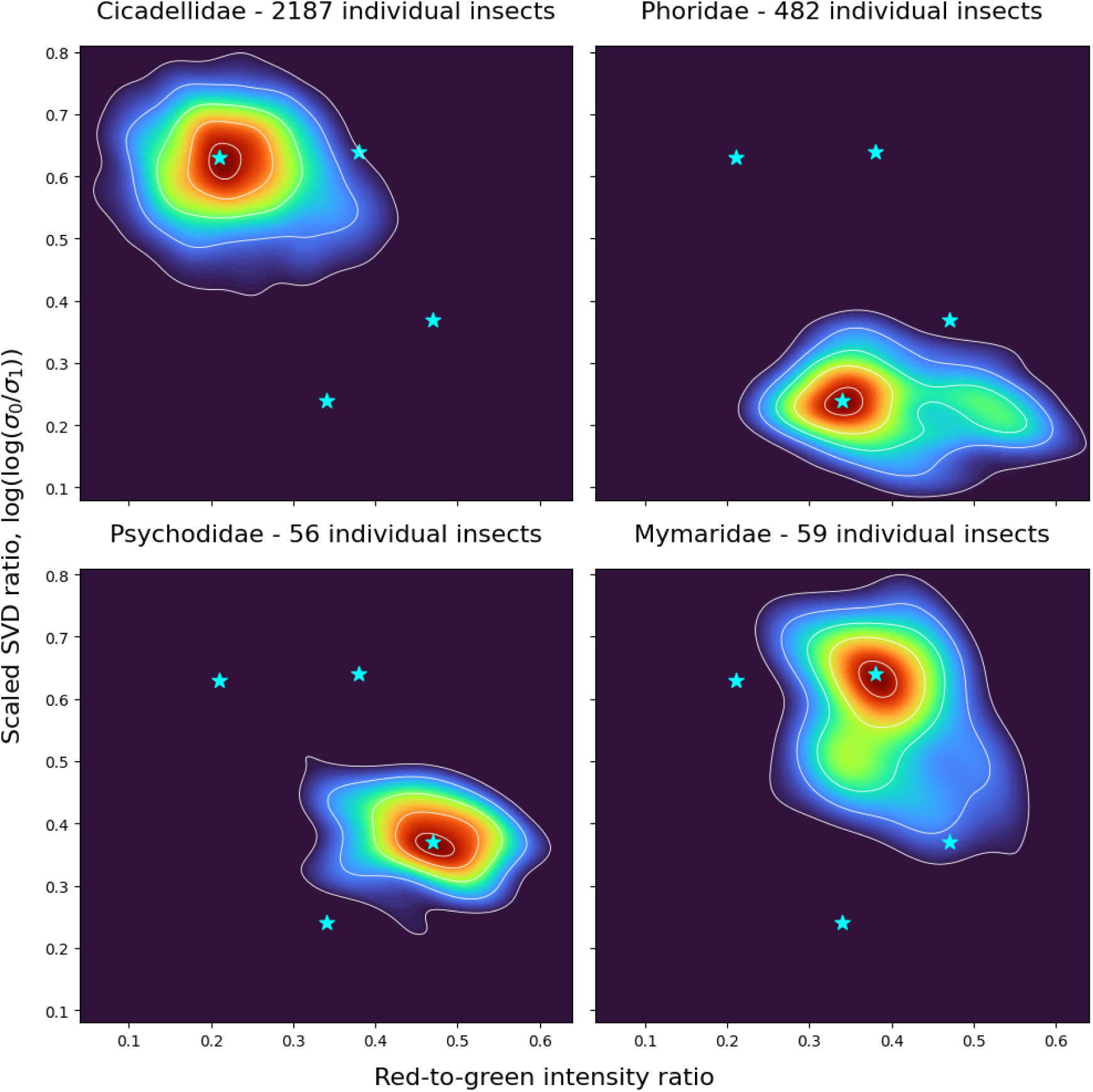
Insect families occupy distinct density distributions in a two-dimensional projection of the five-dimensional feature space used for the estimation of visual diversity. From the outermost to the innermost, the white contour lines enclose 95%, 75%, 50%, 25%, and 10% of the KDE-estimated probability mass, which approximates the expected proportion of individuals. The color scale reflects the point spread (insect images), with warmer colors having a higher concentration of points, cooler colors having a lower density of points. The asterisks depict the four highest densities points.

### 3.2 Sensitivity tests

The performance of GPS-VDI is robust to perturbations in its hyper-parameters. Specifically:

1. The values of the two major hyper-parameters of GPS-VDI, *K* and λ-5 and 40 respectively - were chosen after short experimentation. Figure 8 depicts a more systematic, retrospective, search in the (*K,* λ)parameter space. It demonstrates that the chosen parameters lie within a rather wide performance plateau.
2. A considerable correlation between taxonomic and visual diversity indices does not qualitatively depend on the method used to estimate visual diversity index (Table 2).
3. All five visual features contribute to the utility of GPS-VDI, yet none of them is essential (Table 3).
4. GPS-VDI consistently performs well, when applied to sets of synthetic random traps (Table 4).

**Table 2.** Pearson correlation between manually derived Shannon-Weiber index and VDI. The bolded results correspond to the proposed method. All correlations are significantly different than the null hypothesis of zero correlation (p < 10).

| **Taxonomic level** | **DVI estimation method** | **Pearson Correlation** | **Weighted Pearson correlation** |
|---|---|---|---|
| Order | Grid | 0.80 | 0.81 |
| Order | KDE | 0.72 | 0.75 |
| Order | KDE_loo | 0.67 | 0.66 |
| Family | Grid | 0.66 | 0.74 |
| Family | KDE | 0.53 | 0.65 |
| Family | KDE_loo | 0.47 | 0.48 |

**Table 3:**
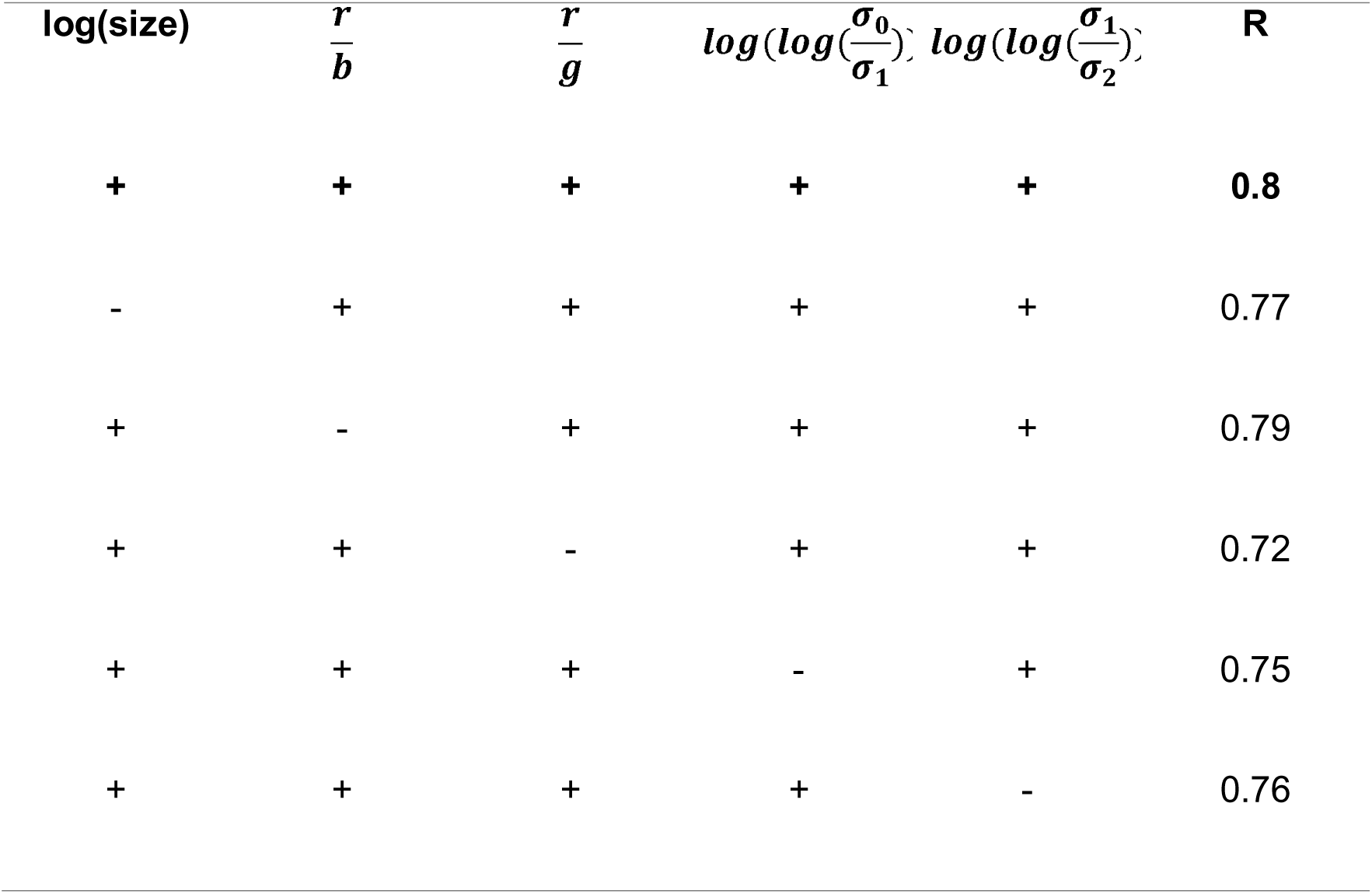
None of the features is essential for performance. The top, right most, value (bold) is the reported performance of GPS-VDI. The five “plus” signs indicate that all five visual features are considered. The following five rows depict four feature combinations and their performances.

**Table 4:** Statistics of 10 sets of random synthetic traps.

|  |  |
|---|---|
| Median | 0.76 |
| Mean | 0.77 |
| Standard deviation | 0.04 |
| Minimum | 0.70 |
| Maximum | 0.84 |

### 3.3 Visual diversity vs. taxonomic diversity

The Shannon index of taxonomic diversity at the order level was strongly and positively correlated with the entropy index of visual diversity (Pearson’s correlation, r = 0.80, p < 0.001). We obtained a similar correlation value when weighting the traps by their insect loads, with higher weights assigned to the denser traps (r = 0.81, Fig. 4). Pearson’s correlation for the Shannon index of taxonomic diversity at the family level was lower (r = 0.67, p < 0.001 for unweighted data, r = 0.74, P<0.001 after using insect abundances as weights, Supplementary Fig. S1).

**Figure 4:**
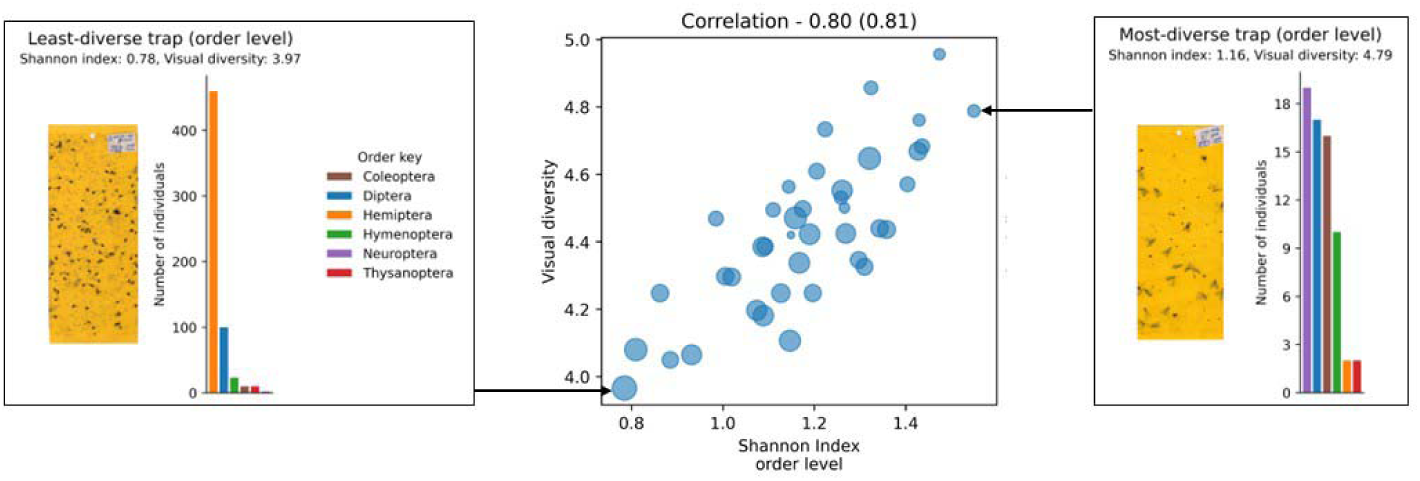
The calculated visual diversity vs. the order-level taxonomic diversity as determined by human experts. Each circle represents a trap (total = 40 traps), the areas of the circles are proportional to the abundance of insects captured. Histograms of two traps illustrating two opposite extreme values are shown: On the left - a trap that received low taxonomic and visual diversity scores, characterized by the dominance of Hemiptera (orange), followed by Diptera (blue). On the right - a trap that received high taxonomic and visual diversity scores, with no dominance of a particular order.

### 3.4 From Computer Vision to Ecological Insight

Regarding our first ecological question, we found that insect abundance was negatively correlated with insect diversity across all 322 sticky traps that comprised our field sample (Pearson’s correlation, r = −0.37, p < 0.001; Fig. 5). As the number of insects captured per trap increased, diversity tended to decrease. Although diversity values exhibited substantial variation among traps with low to moderate insect abundance, traps with the highest insect abundance consistently showed relatively low diversity. The 95% confidence interval for the correlation coefficient (−0.46 to −0.27) indicates a moderate negative association between abundance and diversity. The significant negative correlation between abundance and diversity persisted when we considered the insects caught on all three traps per site as a replicate, i.e computationally pooled each set of three replicates into one large trap (Pearson’s correlation for triplicate: r = −0.39, p < 0.001).

**Figure 5:**
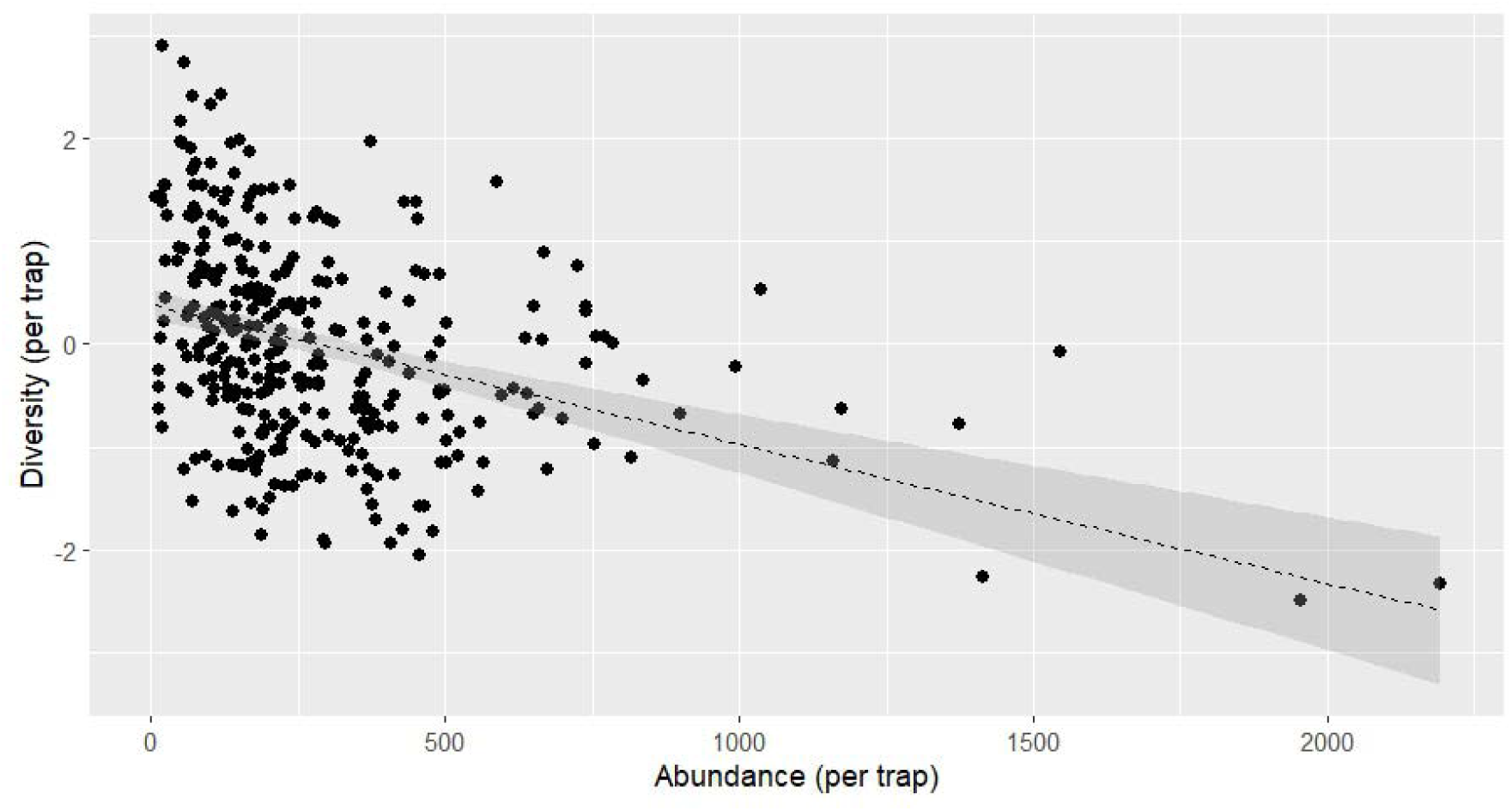
Correlation graph between the visual diversity per trap (y-axis) and the abundance per trap (x-axis). Each point is a trap (total = 322 traps).

We also asked whether diversity and insect abundance differ between habitats (natural vs. agricultural) and seasons (spring vs. summer). A generalized linear mixed-effects model accounting for the paired sampling design revealed significant effects of habitat (F_112.1_=11.1, p < 0.001), season (F_21_=15.3, p < 0.001), and their interaction (F_112.1_=10.2, p = 0.002) on standardized insect diversity (Fig. 6). The significant interaction indicates that seasonal changes in insect diversity differed between habitat types. In natural habitats, mean insect diversity increased from spring to summer, whereas in agricultural habitats the seasonal increase was more moderate, resulting in similar diversity values between habitats during the summer sampling period (GLMM results for triplicate - habitat effects: p = 0.04, season effects: p < 0.001, habitat × season interaction: p = 0.114). The trend was reversed in the effects on insect abundance: In natural habitats, mean insect abundance declined markedly from spring to summer, whereas abundance in agricultural habitats remained relatively stable across seasons. Consequently, insect abundance was substantially higher in natural habitats during spring but converged to similar levels in both habitat types during summer. Accordingly, there was a highly significant habitat × season interaction in insect abundance (χ²=1070.8, p < 0.001), indicating that seasonal changes in abundance differed between agricultural and natural habitats. Habitat also had a significant overall effect (χ²=385.4, p < 0.001), whereas the main effect of season was not significant (χ²=2.206, p = 0.137) (Fig. 7). The GLMM that treated trap triplicates as sampling units generated similar results: habitat effects: p < 0.001, season effects: p = 0.137, habitat × season interaction: p < 0.001).

**Figure 6:**
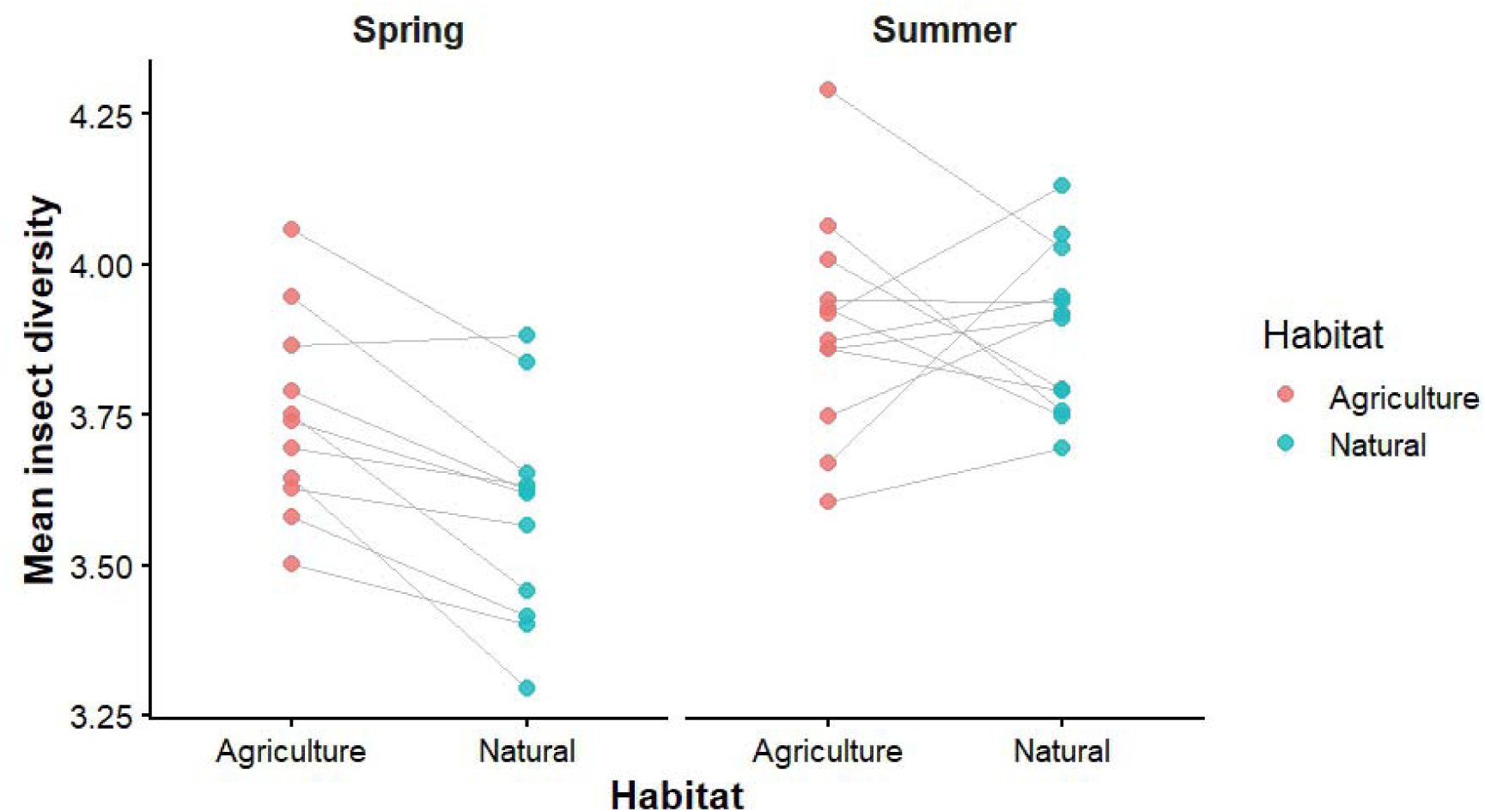
Comparison of mean insect diversity between agricultural and natural habitats during spring and summer. Each point represents the average diversity of three sticky traps within a sampling plot. Grey lines connect corresponding agricultural and natural plots within each sampling pair, illustrating variation in habitat-related differences among sites. The data are presented separately for spring and summer to facilitate comparisons of insect diversity between habitats across seasons.

**Figure 7:**
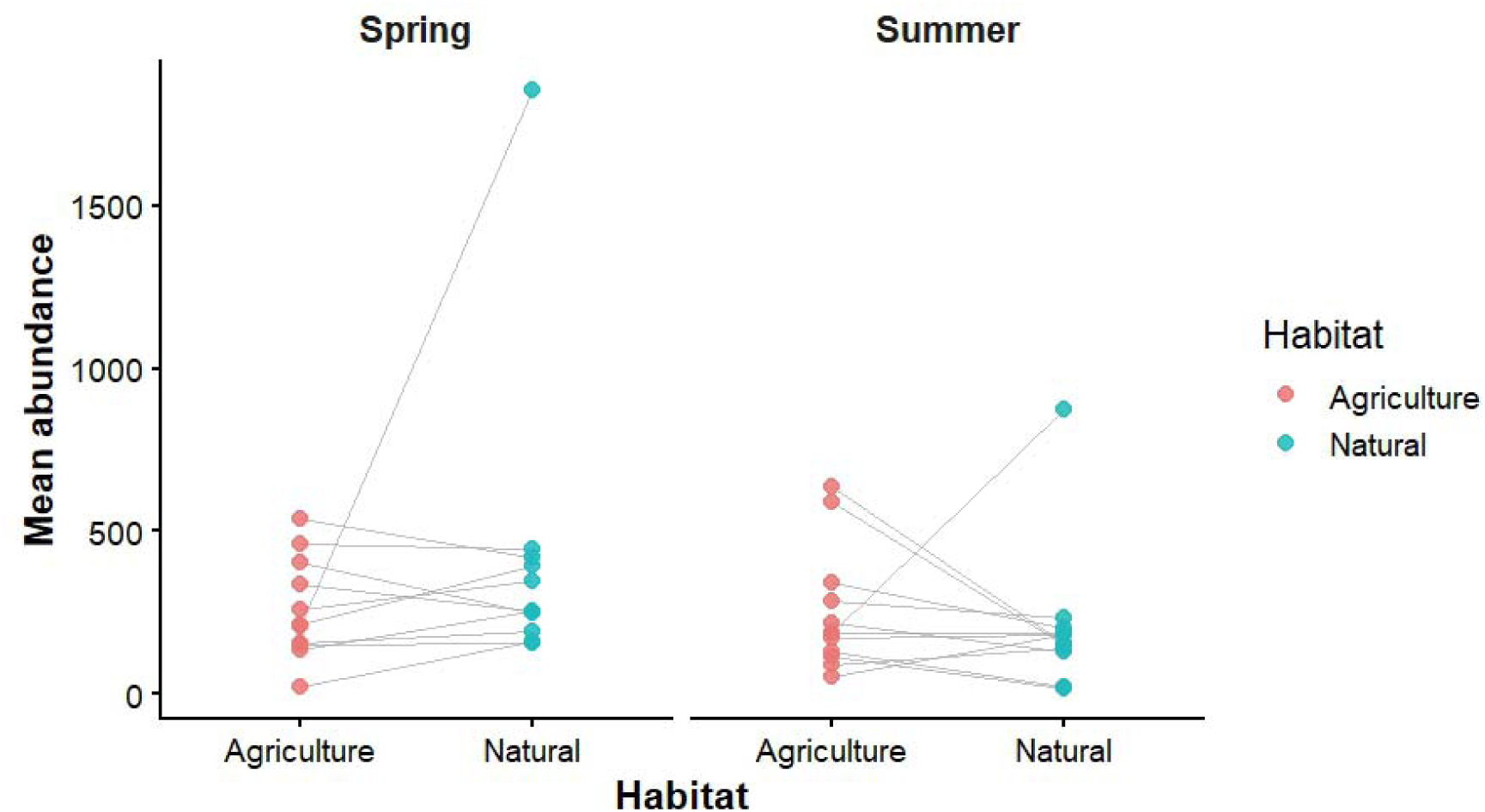
Comparison of mean insect abundance between agricultural and natural habitats during spring and summer. Each point represents the average insect abundance recorded across three sticky traps within a sampling plot. Grey lines connect corresponding agricultural and natural plots within each sampling pair, illustrating variation in habitat-related differences among sites. The data are presented separately for spring and summer to facilitate comparisons of insect abundance between habitats across seasons.

**Figure 8:**
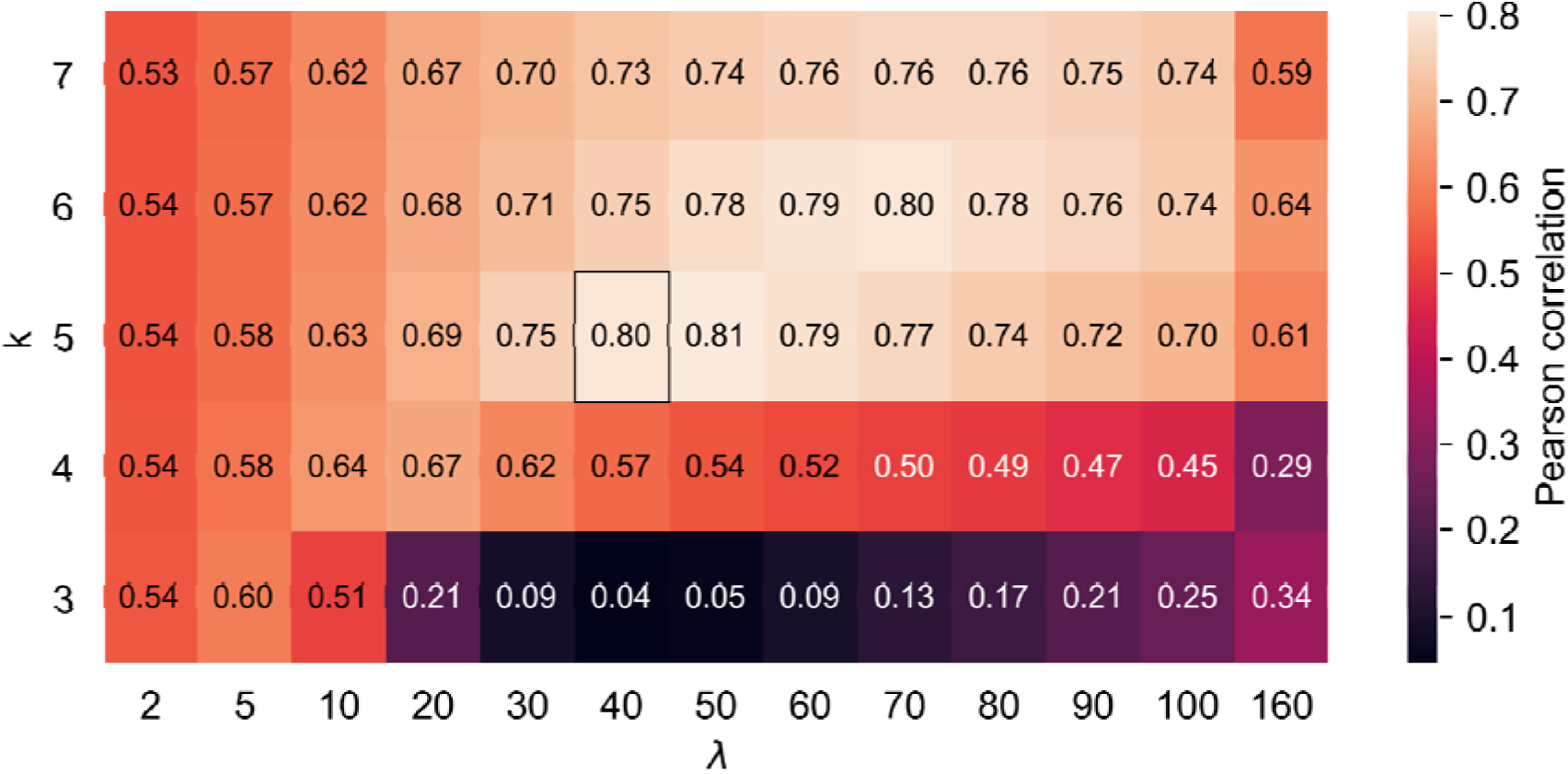
A heat map of correlations between taxonomic and visual diversities. The cells represent different, combinations. The combination used in this study is marked by a black frame.

## 4. Discussion

### 4.1 Concordance between visual diversity and taxonomic diversity

We found a high positive correlation (r=0.80) between the order-level Shannon index of taxonomic diversity and the visual entropy-based index. A positive correlation was also observed at the family-level but was slightly lower (r=0.67). In this case, the correlation improved after using insect abundances as weights (r=0.74). This provides empirical support for our expectation that taxonomic diversity is manifested by visual diversity. This finding is significant because it validates the use of phenotypic traits: body size, shape, and color distribution, as reliable proxies for biological diversity. While many current deep learning models in entomology, based on YOLO and various Convolutional Neural Networks (CNNs), rely on explicit classification into pre-defined categories (order/ family/ species), they often struggle with rare or cryptic species where training data is scarce (Bjerge et al., 2023). By characterizing insects into a five-dimensional hyperspace of engineered features, our model successfully clusters distinct density distributions for different taxonomic groups, such as the clustering observed for Cicadellidae and Psychodidae (Fig. 3). This phenotype-based approach addresses the “taxonomic impediment” - the global shortage of specialists and the extreme labor intensity required to manually identify the world’s vast insect diversity. Our method offers a “rapid estimation” proxy that does not require labeling of training data. It can also be applied to any geographic region or habitat after appropriate validation, because it is not trained on a region-specific insect community, but rather is based on phenotypic characteristics.

### 4.2 Cost-effectiveness of digitized monitoring

Our approach utilizes a “low-tech” capture method (sticky traps and office scanners) combined with “high-tech” tools (image analysis techniques), a workflow that scales effectively for wide-range surveys. This contrasts with more complex autonomous sensor systems like Sticky Pi (Geissmann et al., 2022), Mothbox (Szczygieł et al., 2026a) or the FAIR-Device (Chiavassa et al., 2024). These technologies, while offering high temporal resolution and a rapid output, face challenges regarding power consumption, wireless data transmission, and resilience to harsh environments. Our use of simple traps avoids these constraints and enables comprehensive surveys over wide spatial and temporal ranges at a fraction of the cost of real-time sensor networks (approximately USD 1 for six traps). The computational component, in turn, generates much faster biodiversity estimates than human-based assessments. The Grid Pseudo-Species Visual Diversity Index (GPS-VDI) described here, calculated using image processing alone, saves between 2-4 hours of work on deciphering a single trap by an expert entomologist who identifies the insects to the order or family level. Furthermore, our searchable database of trap images preserves specimen integrity, allowing for retrospective studies, follow-up genetic analyses, and “human-in-the-loop” validation, similar to the Mothbot (Szczygieł et al., 2026b) and BIODISCOVER (Høye et al., 2021) platforms.

### 4.3 Technical Limitations

Along with its advantages, the Grid Pseudo-Species Visual Diversity Index (GPS-VDI) is currently facing several technical limitations common to emerging automated monitoring tools. First, the validation of visual diversity is performed on a single dataset and has not yet been tested across a wide range of habitats. This limitation is echoed by Bjerge et al. (2023), who note that model performance often decreases in different environmental contexts and emphasize the need for combining datasets from diverse locations to achieve better generalization. Second, our visual diversity index does not explicitly reflect functional group diversity, making it difficult to determine ecosystem functions. For example, in agroecosystems one would like to know whether a sample represents a healthy balance of pests, pollinators, and natural enemies. In contrast, models like InsectNet (Chiranjeevi et al., 2024) are trained on labeled images to identify numerous insect species with known ecological roles (e.g., predator, decomposer, or invasive species). Furthermore, the FAIR-Device (Chiavassa et al., 2024) utilizes the AI identification system of the iNaturalist platform to identify insect’s images from Malaise traps. These higher-level taxonomic identifications can be combined with information on the insects’ ecological roles, to establish functional groups that indicate ecosystem integrity. Third, the GPS-VDI is sensitive to high intra-specific variation, particularly sexual dimorphism and stage-specific developmental transformations. In an unsupervised visual feature space, sexually dimorphic individuals of the same species occupy distinct coordinates. This phenotypic divergence can artificially inflate the calculated visual entropy index by treating a single biological species as two distinct visual entities (Chiranjeevi et al., 2024; Schneider et al., 2023; Zhang et al., 2025). This challenge is widely recognized across automated entomological vision systems. For instance, Chiranjeevi et al., (2024) identified sexual dimorphism as a primary obstacle to accurate image-based classification, citing species like the European rhinoceros beetle (*Oryctes nasicornis*), where males possess prominent cephalic horns while females do not. Without prior sex-specific annotations, unsupervised visual diversity estimators inherently absorb this dimorphic variance, underscoring the need for future iterations that calibrate phenotypic hyperspace volume against known biological variance.

A notable logistical limitation of our current workflow is its reliance on flatbed office scanners for trap digitization. Although high-resolution scanning (1,200 dpi) provides standardized illumination and preserves fine morphological details, flatbed scanners are ill-suited for field deployment and require approximately 30 seconds to scan a trap, creating a processing bottleneck in large-scale surveys. To streamline acquisition, recent studies have introduced smartphone-based mobile frameworks (e.g., EyesOnTraps) equipped with real-time focus validation, shadow removal, and perspective correction, enabling rapid in-field capture (Faria et al., 2021; Rosado et al., 2022).

### 4.4 Ecological Insights

In the test of implementing the index, we gained a number of ecological insights.

#### 4.4.1 The Abundance-Diversity relationship

We found a moderate negative correlation (r=−0.37) between insect abundance and visual diversity. Traps with the highest insect loads consistently exhibited the lowest phenotypic heterogeneity. This finding suggests that high-density samples in our study area were often numerically dominated by a few abundant or opportunistic taxa. Indeed, in our study, mosquitoes dominated a high-density trap from a natural habitat near a water canal, and Cicadellidae, known pests of Rosaceae orchards, dominated highly-populated traps from the orchards. This aligns with observations in the Amazon where habitat loss led to a surge in opportunistic “winners” (e.g., mosquitoes) alongside a sharp decline in the overall species richness, a phenomenon known as the “more-individuals” effect (Colares et al., 2025). A similar dynamic is reflected in the work of Kalfas et al. (2023), whose sticky trap surveys in witloof chicory fields were dominated by “critical” pest classes and generalist flies. These species often numbered in the thousands while other taxa remained rare. This abundance-diversity tradeoff highlights the importance of partitioning abundance from biodiversity trends. While total insect abundance is an essential metric for a habitat’s carrying capacity, it can obscure signals of biodiversity loss.

#### 4.4.2 Spatiotemporal monitoring for sustainable landscape management

By capturing biodiversity fluctuations across habitats and seasons, our monitoring provided insights on how seasonal and land-use variation interact to shape insect biodiversity in Mediterranean agroecosystems. We found a significant interaction between habitat type (natural vs. agricultural) and season (spring vs. summer). Natural habitats exhibited a sharper seasonal increase in insect diversity than paired agricultural habitats. This relative stability in agricultural habitats likely reflects the influence of continuous irrigation, which buffers against the extreme seasonal drying characteristic of Mediterranean summers. Furthermore, the agricultural sites in this study featured tree species with stable foliage that does not change between spring and summer, providing consistent structural resources and microclimates. This stability of plant cover is a known driver of arthropod abundance and diversity (Litovska et al., 2025). In contrast, the more pronounced fluctuations in natural habitats are often tied to seasonal resource pulses; for example, Segoli et al. (2020) demonstrated that parasitoid populations are most abundant in natural desert habitats following spring rains but shift toward irrigated vineyards during the dry summer months as natural vegetation desiccates. Surprisingly, our results showed that diversity values were lower in natural areas compared to agricultural areas. This finding differs from the prevailing literature, which typically identifies semi-natural habitats as essential refuges for biodiversity within agroecosystems (Litovska et al., 2025). For instance, Shapira et al. (2018) found that parasitoid diversity was highest in natural habitats and decreased toward the center of vineyards. This discrepancy may reflect the fact that we, as well as Shapira et al. (2018), sampled the plots only 2-3 times per season. Such sparse sampling may have missed population dynamics at high temporal resolutions. The benefits of high-resolution monitoring were illustrated by Kalfas et al. (2023), who showed that automated sticky trap analysis can accurately track the weekly dynamics of pest species. This provides the data necessary for “timely decision-making” in sustainable landscape management. Similarly, Geissman et al. (2022) monitored insects automatically at an even finer temporal scale to characterize their circadian activity schedules.

### 4.5 Future Directions and Global Policy Implications

The Grid Pseudo-Species Visual Diversity Index (GPS-VDI) developed in this study offers a powerful, cost-effective complementation to traditional monitoring, which remains severely constrained by high logistical costs and a global shortage of taxonomic specialists (Preti et al., 2021).

Crucially, automated visual proxies are not intended to replace human taxonomists, but rather to expand their capacity by delegating the labor-intensive, routine screening of bulk trap samples to AI-driven algorithms. Beyond easing this “taxonomic bottleneck”, GPS-VDI provides a rapid assessment proxy for evaluating the ecological value of natural and agricultural landscapes. Its low cost and technical simplicity make it an ideal tool for standardized, long-term monitoring to examine the effects of diverse land management practices. For example, GPS-VDI is a promising tool to monitor the long-term effects of agro-voltaic systems, a relatively new practice, on insect communities. These effects have not yet been fully explored (Walston et al., 2024).

This method represents a universal tool capable of providing standardized metrics for global insect populations, as it can be applied to any habitat and region in the world. Thus, it can be integrated into a future toolkit for global biodiversity assessment. Furthermore, this visual diversity assessment provides a fundamental component within an existing set of methods. For instance, image-based identification of insects can be effectively integrated with DNA metabarcoding. While the visual index captures rapid shifts in abundance and phenotypic heterogeneity, metabarcoding can resolve the species composition of cryptic or undescribed taxa from the same trapped specimens (Fujisawa et al., 2023). By integrating these high-throughput “big data” strategies, the scientific community can build a scalable, standardized framework to combat global insect declines and ensure that insect biodiversity is accurately represented, alongside other biodiversity assessments, in conservation and policy decisions (Szczygieł et al., 2026b).

Beyond local ecological monitoring, GPS-VDI directly supports international conservation policy. Under the Convention on Biological Diversity (CBD) Kunming-Montreal Global Biodiversity Framework (GBF), signatory nations require continuous, standardized monitoring systems to track progress toward global biodiversity targets and update National Biodiversity Strategies and Action Plans (NBSAPs). Major geographic, temporal, and taxonomic data gaps persist for insects due to traditional sorting bottlenecks. By providing a standardized, “human-out-of-the-loop” visual diversity metric integrated into the open-access, web-based STARdbi database, users worldwide can upload trap imagery, continually expand the repository, and rapidly receive automated GPS-VDI scores for their samples. GPS-VDI generates primary data that can be readily translated into Essential Biodiversity Variables (EBVs) and scalable indicator toolkits. This interoperable workflow aligns with the CBD’s vision for global biodiversity observation networks (e.g., GBiOS) operating under FAIR data principles, providing land managers and decision-makers with the scalable evidence needed for adaptive landscape planning (Gonzalez et al., 2022).

## 5. Conclusions

This study demonstrates that flying insect biodiversity is fundamentally reflected in the morphological heterogeneity of insect bodies, validating the Grid Pseudo-species Visual Diversity Index (GPS-VDI) as a reliable visual-entropy proxy for taxonomic diversity. By establishing strong concordance with expert-derived Shannon-Weaver indices at both the order and family levels, this phenotype-based approach circumvents the traditional “taxonomic impediment” without requiring pre-labeled training datasets, region-specific classifiers, or destructive specimen processing. When applied across field environments, GPS-VDI effectively captured key ecological dynamics, revealing a moderate inverse relationship between insect abundance and visual diversity driven by the numerical dominance of opportunistic taxa, as well as distinct spatiotemporal microclimatic buffering between agricultural and natural habitats. Designed to expand the operational capacity of taxonomists rather than replace them, this workflow delegates routine bulk-sample screening to automated algorithms while preserving digital images in the open-access STARdbi database for community data expansion and rapid automated scoring. Ultimately, by combining low-cost sticky traps and flatbed scanning with automated visual feature extraction, GPS-VDI offers an accessible, field-ready framework that integrates with molecular methods like DNA metabarcoding and delivers standardized primary data aligned with Essential Biodiversity Variables (EBVs) and the Global Biodiversity Framework.

## Acknowledgements

This study was supported by the Israel Ministry of Energy (grant #221-11-041) and by the Israel Council of Higher Education (Research Center for Biodiversity in Agroecosystems). LCL was supported by PhD scholarships from the University of Haifa and the Jewish National Fund (KKL).

Special thanks to the expert taxonomists-Ronen Shapir, Nitsan Nachtom Catalan, Levona Bodner and Elizabeth (Liz) Morgulis. Michael Yair - STARdbi Software Developer for technical support, and to all farmers - for their cooperation.

## Declaration of generative AI and AI-assisted technologies in the manuscript preparation process

During the preparation of this work, the authors used ChatGPT and NotebookLM to improve and proof-read the text, organize literary sources and write code. After using these tools, the authors reviewed and edited the content as needed and take full responsibility for the content of the published article.

## Data availability

All trap images files, data, and analysis scripts are available at DOI: 10.17632/6pbgs93kdf and Python code at https://github.com/chenKeasar/gps-vdi.

## Appendix Supplementary data

**Figure S1:**
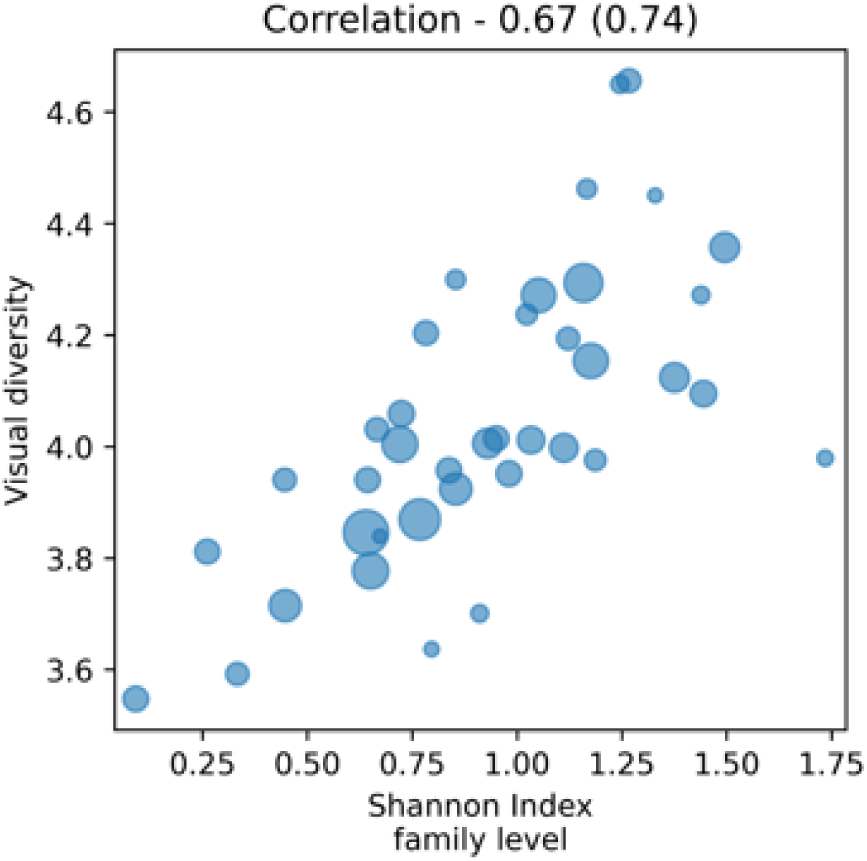
The calculated visual diversity vs. the family-level taxonomic diversity as determined by human experts. Each circle represents a trap (total = 40 traps), the areas of the circles are proportional to the abundance of insects captured.

---

**Table S1:** Authoritative taxonomic identifiers for every taxon using the Integrated Taxonomic Information System (ITIS TSN). This table presents the identifiers for all insect orders (bold) and families identified in the sticky traps by expert taxonomists.

| Taxon | ITIS TSN |
|---|---|
| ***Araneae*** | 82732 |
| ***Diptera*** | 118831 |
| *Dolichopodidae* | 136824 |
| *Syrphidae* | 139621 |
| *Phoridae* | 138921 |
| *Chloropidae* | 147644 |
| *Sepsidae* | 144975 |
| *Lonchaeidae* | 142167 |
| *Tephritidae* | 142604 |
| *Muscidae* | 150025 |
| *Anthomyiidae* | 148685 |
| *Tachinidae* | 152643 |
| *Calliphoridae* | 151495 |
| *Psychodidae* | 125351 |
| ***Coleoptera*** | 109216 |
| *Coccinellidae* | 114329 |
| ***Hemiptera*** | 103359 |
| *Lygaeidae* | 107674 |
| *Cicadellidae* | 109171 |
| ***Hymenoptera*** | 152741 |
| *Formicidae* | 154193 |
| *Ceraphronidae* | 154066 |
| *Encyrtidae* | 153945 |
| *Eulophidae* | 153770 |
| *Mymaridae* | 153659 |
| *Trichogrammatidae* | 153728 |
| *Platygastridae* | 709288 |
| *Scelionidae* | 154068 |
| ***Lepidoptera*** | 117232 |
| ***Neuroptera*** | 114999 |
| *Chrysopidae* | 115081 |
| *Berothidae* | 115083 |
| *Mantispidae* | 115079 |
| ***Thysanoptera*** | 103348 |
| ***Trombidiformes*** | 82769 |

## References

Bates, D., Mächler, M., Bolker, B., Walker, S., 2015. Fitting Linear Mixed-Effects Models Using lme4. J. Stat. Soft. 67. 10.18637/jss.v067.i01

Billeter, R., Liira, J., Bailey, D., Bugter, R., Arens, P., Augenstein, I., Aviron, S., Baudry, J., Bukacek, R., Burel, F., Cerny, M., De Blust, G., De Cock, R., Diekötter, T., Dietz, H., Dirksen, J., Dormann, C., Durka, W., Frenzel, M., Hamersky, R., Hendrickx, F., Herzog, F., Klotz, S., Koolstra, B., Lausch, A., Le Coeur, D., Maelfait, J.P., Opdam, P., Roubalova, M., Schermann, A., Schermann, N., Schmidt, T., Schweiger, O., Smulders, M.J.M., Speelmans, M., Simova, P., Verboom, J., Van Wingerden, W.K.R.E., Zobel, M., Edwards, P.J., 2007. Indicators for biodiversity in agricultural landscapes: a pan-European study: Biodiversity in European Agro-ecosystems. Journal of Applied Ecology 45, 141–150. 10.1111/j.1365-2664.2007.01393.x

Biswas, D., Tiwari, A., 2024. Utilizing Computer Vision and Deep Learning to Detect and Monitor Insects in Real Time by Analyzing Camera Trap Images. Natural and Engineering Sciences 9, 280–292. 10.28978/nesciences.1575480

Bjerge, K., Alison, J., Dyrmann, M., Frigaard, C.E., Mann, H.M.R., Høye, T.T., 2023. Accurate detection and identification of insects from camera trap images with deep learning. PLOS Sustain Transform 2, e0000051. 10.1371/journal.pstr.0000051

Buchner, D., Sinclair, J.S., Ayasse, M., Beermann, A.J., Buse, J., Dziock, F., Enss, J., Frenzel, M., Hörren, T., Li, Y., Monaghan, M.T., Morkel, C., Müller, J., Pauls, S.U., Richter, R., Scharnweber, T., Sorg, M., Stoll, S., Twietmeyer, S., Weisser, W.W., Wiggering, B., Wilmking, M., Zotz, G., Gessner, M.O., Haase, P., Leese, F., 2025. Upscaling biodiversity monitoring: Metabarcoding estimates 31,846 insect species from Malaise traps across Germany. Molecular Ecology Resources 25, e14023. 10.1111/1755-0998.14023

Burger, J., 2006. Bioindicators: Types, Development, and Use in Ecological Assessment and Research. Environmental Bioindicators 1, 22–39. 10.1080/15555270590966483

Chiavassa, J.A., Kraft, M., Noack, P., Walther, S., Kirse, A., Scherber, C., 2024. The Field Automatic Insect Recognition-Device—A Non-Lethal Semi-Automatic Malaise Trap for Insect Biodiversity Monitoring: Proof of Concept. Ecology and Evolution 14, e70642. 10.1002/ece3.70642

Chiranjeevi, S., Saadati, M., Deng, Z.K., Koushik, J., Jubery, T.Z., Mueller, D.S., O’Neal, M., Merchant, N., Singh, Aarti, Singh, A.K., Sarkar, S., Singh, Arti, Ganapathysubramanian, B., 2024. InsectNet: Real-time identification of insects using an end-to-end machine learning pipeline. PNAS Nexus 4, pgae575. 10.1093/pnasnexus/pgae575

Ciampi, L., Zeni, V., Incrocci, L., Canale, A., Benelli, G., Falchi, F., Amato, G., Chessa, S., 2023. A deep learning-based pipeline for whitefly pest abundance estimation on chromotropic sticky traps. Ecological Informatics 78, 102384. 10.1016/j.ecoinf.2023.102384

Colares, L.F., Peres, C.A., Dambros, C.S., 2025. Life history induces markedly divergent insect responses to habitat loss. Journal of Animal Ecology 1365–2656.70117. 10.1111/1365-2656.70117

Duelli, P., Obrist, M.K., 2003. Biodiversity indicators: the choice of values and measures. Agriculture, Ecosystems & Environment 98, 87–98. 10.1016/S0167-8809(03)00072-0

Epanechnikov, V.A., 1969. Non-parametric estimation of a multidimensional probability density. Theory of Probability \& Its Applications 14, 153–158.

Fujisawa, T., Noguerales, V., Meramveliotakis, E., Papadopoulou, A., Vogler, A.P., 2023. Image-based taxonomic classification of bulk insect biodiversity samples using deep learning and domain adaptation. Systematic Entomology 48, 387–401. 10.1111/syen.12583

Galler, C., Von Haaren, C., Albert, C., 2015. Optimizing environmental measures for landscape multifunctionality: Effectiveness, efficiency and recommendations for agri-environmental programs. Journal of Environmental Management 151, 243–257. 10.1016/j.jenvman.2014.12.011

Gao, Y., Xue, X., Qin, G., Li, K., Liu, J., Zhang, Y., Li, X., 2024. Application of machine learning in automatic image identification of insects - a review. Ecological Informatics 80, 102539. 10.1016/j.ecoinf.2024.102539

Geissmann, Q., Abram, P.K., Wu, D., Haney, C.H., Carrillo, J., 2022. Sticky Pi is a high-frequency smart trap that enables the study of insect circadian activity under natural conditions. PLoS Biol 20, e3001689. 10.1371/journal.pbio.3001689

Gramfort, A., Papadopoulo, T., Olivi, E., Clerc, M., 2011. Forward Field Computation with OpenMEEG. Computational Intelligence and Neuroscience 2011, 1–13. 10.1155/2011/923703

Harris, C.R., Millman, K.J., Van Der Walt, S.J., Gommers, R., Virtanen, P., Cournapeau, D., Wieser, E., Taylor, J., Berg, S., Smith, N.J., Kern, R., Picus, M., Hoyer, S., Van Kerkwijk, M.H., Brett, M., Haldane, A., Del Río, J.F., Wiebe, M., Peterson, P., Gérard-Marchant, P., Sheppard, K., Reddy, T., Weckesser, W., Abbasi, H., Gohlke, C., Oliphant, T.E., 2020. Array programming with NumPy. Nature 585, 357–362. 10.1038/s41586-020-2649-2

Høye, T.T., Ärje, J., Bjerge, K., Hansen, O.L.P., Iosifidis, A., Leese, F., Mann, H.M.R., Meissner, K., Melvad, C., Raitoharju, J., 2021. Deep learning and computer vision will transform entomology. Proc. Natl. Acad. Sci. U.S.A. 118, e2002545117. 10.1073/pnas.2002545117

Hunter, J.D., 2007. Matplotlib: A 2D Graphics Environment. Comput. Sci. Eng. 9, 90–95. 10.1109/MCSE.2007.55

IPBES, 2019. Summary for policymakers of the global assessment report on biodiversity and ecosystem services. Zenodo. 10.5281/ZENODO.3553579

J. S., R., E., V.M., Reddy, G.R., Akhil, V.S., N., R., D., S., 2024. DNA Barcoding: Accelerating Insect Species Discovery and Biodiversity Documentation. J. Adv. Biol. Biotechnol. 27, 709–720. 10.9734/jabb/2024/v27i71030

Kalfas, I., De Ketelaere, B., Bunkens, K., Saeys, W., 2023. Towards automatic insect monitoring on witloof chicory fields using sticky plate image analysis. Ecological Informatics 75, 102037. 10.1016/j.ecoinf.2023.102037

Keasar, T., Yair, M., Gottlieb, D., Cabra-Leykin, L., Keasar, C., 2024. STARdbi: A pipeline and database for insect monitoring based on automated image analysis. Ecological Informatics 80, 102521. 10.1016/j.ecoinf.2024.102521

Kent, M., Smart, N., 1981. A method for habitat assessment in agricultural landscapes. Applied Geography 1, 9–30. 10.1016/0143-6228(81)90003-5

Kittichai, V., Pengsakul, T., Chumchuen, K., Samung, Y., Sriwichai, P., Phatthamolrat, N., Tongloy, T., Jaksukam, K., Chuwongin, S., Boonsang, S., 2021. Deep learning approaches for challenging species and gender identification of mosquito vectors. Sci Rep 11, 4838. 10.1038/s41598-021-84219-4

Kuznetsova, A., Brockhoff, P.B., Christensen, R.H.B., 2017. lmerTest Package: Tests in Linear Mixed Effects Models. J. Stat. Soft. 82. 10.18637/jss.v082.i13

Litovska, I., Van Der Plas, F., Kleijn, D., 2025. Arthropod abundance is most strongly driven by crop and semi-natural habitat type rather than management in an intensive agricultural landscape in the Netherlands. Agriculture, Ecosystems & Environment 378, 109298. 10.1016/j.agee.2024.109298

Long, P.R., Benz, D., Martin, A.C., Holland, P.W.A., Macias-Fauria, M., Seddon, A.W.R., Hagemann, R., Frost, T.K., Simpson, A., Power, D.J., Slaymaker, M.A., Willis, K.J., 2018. LEFT —A web-based tool for the remote measurement and estimation of ecological value across global landscapes. Methods Ecol Evol 9, 571–579. 10.1111/2041-210X.12924

McKinney, W., 2010. Data structures for statistical computing in Python. scipy.

Mörtberg, U.M., Balfors, B., Knol, W.C., 2007. Landscape ecological assessment: A tool for integrating biodiversity issues in strategic environmental assessment and planning. Journal of Environmental Management 82, 457–470. 10.1016/j.jenvman.2006.01.005

Parzen, E., 1962. On estimation of a probability density function and mode. The Annals of Mathematical Statistics 33, 1065–1076.

Pinto-Ledezma, J.N., Schweiger, A.K., Guzmán Q., J.A., Cavender-Bares, J., 2025. Plant diversity across dimensions: Coupling biodiversity measures from the ground and the sky. Sci. Adv. 11, eadr0278. 10.1126/sciadv.adr0278

Plant, C., Thai, S.M., Shao, J., Theis, F.J., Meyer-Baese, A., Böhm, C., 2012. Measuring Non-Gaussianity by Phi-Transformed and Fuzzy Histograms. Advances in Artificial Neural Systems 2012, 1–13. 10.1155/2012/962105

Preti, M., Verheggen, F., Angeli, S., 2021. Insect pest monitoring with camera-equipped traps: strengths and limitations. J Pest Sci 94, 203–217. 10.1007/s10340-020-01309-4

R Core Team, 2022. R: A language and environment for statistical computing, Vienna, Austria. URL https://www.R-project.org/.

Ren, S., He, K., Girshick, R., Sun, J., 2015. Faster R-CNN: Towards Real-Time Object Detection with Region Proposal Networks, in: Cortes, C., Lawrence, N., Lee, D., Sugiyama, M., Garnett, R. (Eds.), Advances in Neural Information Processing Systems. Curran Associates, Inc.

Reznik, Y.A., Chandrasekhar, V., Takacs, G., Chen, D.M., Tsai, S.S., Girod, B., 2010. Fast quantization and matching of histogram-based image features, in: Tescher, A.G. (Ed.),. Presented at the Proc. SPIE 7798, Applications of Digital Image Processing XXXIII, pp. 154–167. 10.1117/12.862362

Rosenblatt, M., 1956. Remarks on some nonparametric estimates of a density function. The Annals of Mathematical Statistics 27, 832–837.

Schneider, S., Taylor, G.W., Kremer, S.C., Burgess, P., McGroarty, J., Mitsui, K., Zhuang, A., deWaard, J.R., Fryxell, J.M., 2022. Bulk arthropod abundance, biomass and diversity estimation using deep learning for computer vision. Methods Ecol Evol 13, 346–357. 10.1111/2041-210X.13769

Schneider, S., Taylor, G.W., Kremer, S.C., Fryxell, J.M., 2023. Getting the bugs out of AI: Advancing ecological research on arthropods through computer vision. Ecology Letters 26, 1247–1258. 10.1111/ele.14239

Segoli, M., Kishinevsky, M., Rozenberg, T., Hoffmann, I., 2020. Parasitoid Abundance and Community Composition in Desert Vineyards and Their Adjacent Natural Habitats. Insects 11, 580. 10.3390/insects11090580

Seyidbayli, C., Fengler, B., Szafranski, D., Reinhardt, A., 2025. Acoustic Trap Design for Biodiversity Detection. IoT 6, 58. 10.3390/iot6040058

Shapira, I., Gavish-Regev, E., Sharon, R., Harari, A.R., Kishinevsky, M., Keasar, T., 2018. Habitat use by crop pests and natural enemies in a Mediterranean vineyard agroecosystem. Agriculture, Ecosystems & Environment 267, 109–118. 10.1016/j.agee.2018.08.012

Stephenson, P., 2020. Technological advances in biodiversity monitoring: applicability, opportunities and challenges. Current Opinion in Environmental Sustainability 45, 36–41. 10.1016/j.cosust.2020.08.005

Szczygieł, Hubert A., Johns, B., Fortet, B., Dent, D.H., Quitmeyer, K., Quitmeyer, A., 2026a. Mothbox and Mothbot: Automated light trap and data processing system for scalable insect monitoring. Methods Ecol Evol 17, 2082–2091. 10.1111/2041-210x.70327

Szczygieł, Hubert A, Quitmeyer, A., Johns, B., Dent, D.H., Sondhi, Y., Zuleta, D., Antonelli, A., 2026b. Simultaneous Automated Insect Monitoring Across a Remote Tropical Elevation Gradient With Mothbox. Integrative And Comparative Biology 66, icag072. 10.1093/icb/icag072

Teixeira, A.C., Ribeiro, J., Morais, R., Sousa, J.J., Cunha, A., 2023. A Systematic Review on Automatic Insect Detection Using Deep Learning. Agriculture 13, 713. 10.3390/agriculture13030713

Tielens, E.K., Stepanian, P.M., Kelly, J.F., 2025. Systematic Continental Scale Monitoring by Weather Surveillance Radar Shows Fewer Insects Above Warming Landscapes in the United States. Global Change Biology 31, e70587. 10.1111/gcb.70587

Van Klink, R., Sheard, J.K., Høye, T.T., Roslin, T., Do Nascimento, L.A., Bauer, S., 2024. Towards a toolkit for global insect biodiversity monitoring. Phil. Trans. R. Soc. B 379, 20230101. 10.1098/rstb.2023.0101

Walston, L.J., Hartmann, H.M., Fox, L., Macknick, J., McCall, J., Janski, J., Jenkins, L., 2024. If you build it, will they come? Insect community responses to habitat establishment at solar energy facilities in Minnesota, USA. Environ. Res. Lett. 19, 014053. 10.1088/1748-9326/ad0f72

Waskom, M., 2021. seaborn: statistical data visualization. JOSS 6, 3021. 10.21105/joss.03021

Wathern, P., Young, S.N., Brown, I.W., Roberts, D.A., 1986. Ecological evaluation techniques. Landscape Planning 12, 403–420. 10.1016/0304-3924(86)90005-5

Wu, Y., Kirillov, A., Massa, F., Lo, W.-Y., Girshick, R., 2019. Detectron2. Facebook AI Research.

Zhang, L., Sun, S., Zhao, H., Li, Z., Li, D., 2025. Multi-species insect recognition method based on computer visions: Sustainable agricultural development. Ecological Informatics 88, 103125. 10.1016/j.ecoinf.2025.103125

Zou, Z., Chen, K., Shi, Z., Guo, Y., Ye, J., 2023. Object Detection in 20 Years: A Survey. Proc. IEEE 111, 257–276. 10.1109/JPROC.2023.3238524

